# A Fully Automated, Data-Driven Approach for Dimensionality Reduction and Clustering in Single-Cell RNA-seq Analysis

**DOI:** 10.1101/2025.10.06.680609

**Authors:** Hyun Kim, Faeyza Rishad Ardi, Kévin Spinicci, Jae Kyoung Kim

**Affiliations:** Biomedical Mathematics Group, Pioneer Research Center for Mathematical and Computational Sciences, Institute for Basic Science, Daejeon 34126, Republic of Korea; School of Computing, KAIST, Daejeon 34141, Republic of Korea; Department of Mathematical Sciences, KAIST, Daejeon 34141, Republic of Korea; Department of Medicine, College of Medicine, Korea University, Seoul, 02841, Republic of Korea

**Keywords:** Clustering analysis, Dimensionality reduction, scRNA-seq, Cell type identification, Automated pipeline

## Abstract

Single-cell RNA sequencing (scRNA-seq) provides deep insights into cellular heterogeneity but demands robust dimensionality reduction (DR) and clustering to handle high-dimensional, noisy data. Many DR and clustering approaches rely on user-defined parameters, undermining reliability. Even automated clustering methods like ChooseR and MultiK still employ fixed principal component defaults, limiting their full automation. To overcome this limitation, we propose a fully automated clustering approach by integrating scLENS—a method for optimal PC selection—with these tools. Our fully automated approach improves clustering performance by ∼14% for ChooseR and ∼10% for MultiK and identifies additional cell subtypes, highlighting the advantages of adaptive, data-driven DR.

**Highlights:**

- Fully automated, data-driven clustering pipeline for scRNA-seq analysis.
- Addresses the limitation of fixed principal component defaults in DR.
- Data-driven pipeline improves clustering performance by 10-14%.
- Performance gains are most pronounced on high-sparsity, high-skewness data.

**Author Summary:** A fundamental goal in modern biology is to create detailed cellular maps of complex tissues to understand their function in health and disease. Achieving this level of detail is now possible through single-cell sequencing, a technology that generates massive, high-dimensional, and noisy datasets of individual cells’ genetic profiles. However, accurately identifying cell-types from these datasets remains a major analytical hurdle, as current computational methods rely on subjective parameters that compromise reliability and reproducibility.

To address this challenge, we developed a fully automated pipeline that integrates scLENS, an adaptive noise filtering method, with automated cell grouping tools. By providing optimal parameters for noise filtering and cell grouping, our pipeline eliminates their reliance on subjective, fixed parameters, thereby significantly improving accuracy in cell-type classification. This tool, characterized by its high clustering efficiency, can accelerate discoveries in fields like cancer biology and immunology by providing a robust and reproducible analytical platform.

## Introduction

Many recent studies utilize single-cell RNA sequencing (scRNA-seq) because it provides extensive transcriptomic information[1,2]. Most scRNA-seq analysis pipelines begin by filtering out low-quality cells and genes, followed by normalization to reduce distribution skewness, and, if necessary, feature selection or batch correction before dimension reduction (DR) [2–5]. This DR step mitigates the curse of dimensionality by extracting only biologically meaningful information, and the resulting lower-dimensional embedding is then used for clustering [2–4,6]. After these preliminary analyses, various downstream steps become relevant, including differential expression analysis (DEA) to identify cell functions, trajectory analysis to trace cell lineage, and pseudotime analysis to capture cell differentiation over time [7–11]. Because both DR and clustering directly influence these follow-up methods, they are regarded as indispensable processes in the scRNA-seq workflow [2,4,6].

In practice, DR is performed via either linear methods like principal component analysis (PCA) or nonlinear methods such as t-SNE, UMAP, and variational autoencoders (VAE). PCA, in particular, remains a popular tool due to its computational efficiency and interpretability [12–19].

Despite its utility, misselecting the number of principal components (PCs) during the PCA process can critically compromise the embedding quality [20]. Given this significance, many studies still rely on default software settings or employ naïve criteria based on the dataset’s total number of cells [21–23]. To address this issue, several approaches—including Elbow plots, variance-based criteria, and Horn’s parallel analysis—have been proposed [24–26]. However, each still depends partly on user decisions or has scalability issues. As a result, many widely used packages, such as Seurat and Scanpy, continue to provide a fixed number of PCs as their default [17,18]. Recently, however, scLENS emerged as a fast and accurate solution for automatically determining the optimal number of PCs [27]. This prompts the question of whether replacing the default PCA settings with scLENS’s data-driven approach could enhance the performance of existing pipelines.

In this study, we sought to answer that question by reimplementing scLENS—originally developed in Julia—into a Python environment, where numerous analysis packages are widely used [28]. This Python-based scLENS was combined with two data-driven clustering algorithms, ChooseR and MultiK, both of which can automatically derive an optimal number of clusters via consensus clustering (Fig. 1a-c; see details in Methods) [29,30]. However, the original algorithms for these approaches rely on fixed PC defaults in the DR step (*N* = 30 for MultiK and *N* = 100 for ChooseR; Fig. 1a-c), regardless of the biological complexity of the data [29,30].

**Figure 1.**
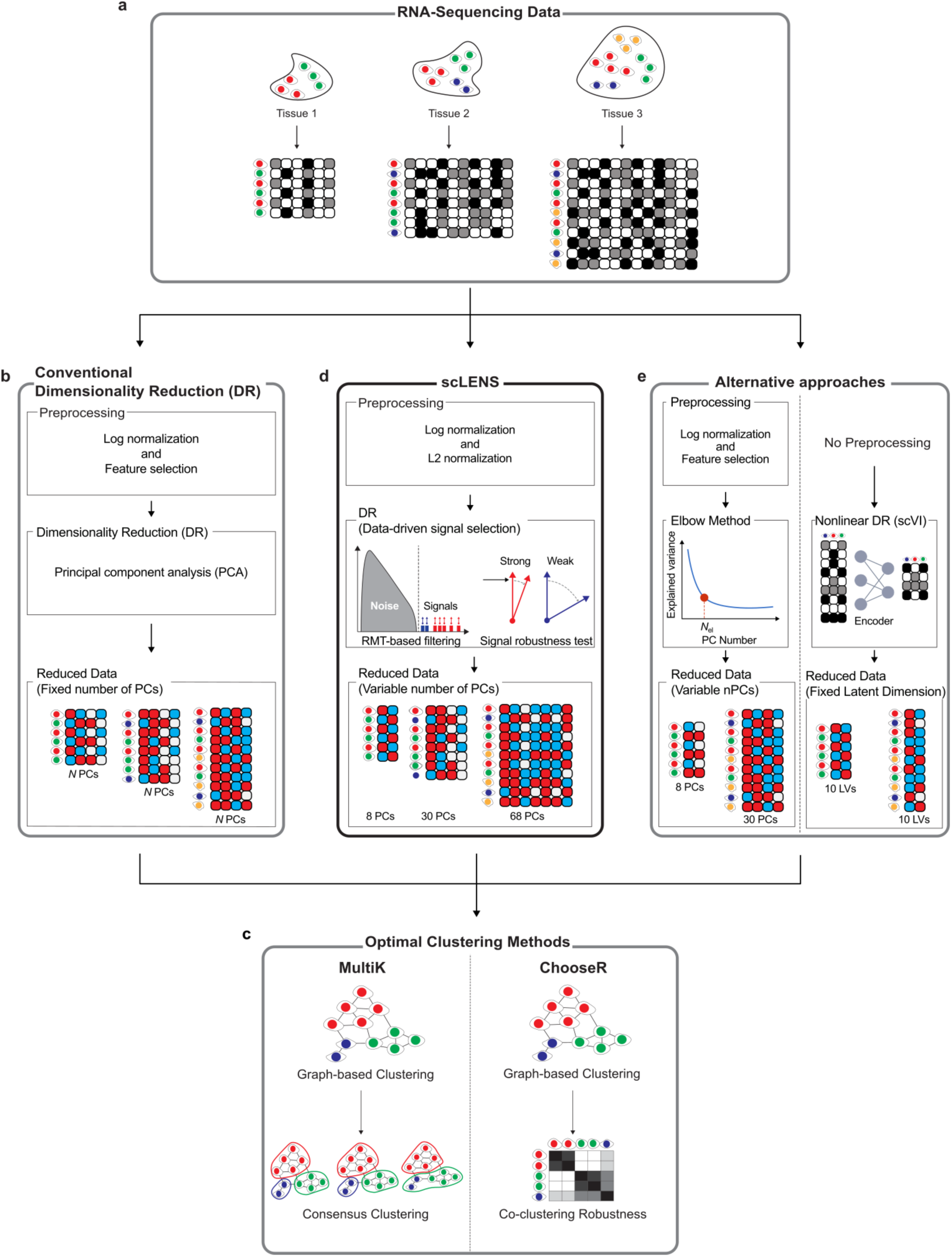
The dimensionality reduction (DR) step in conventional clustering workflows is replaced with scLENS. **a**. Single-cell RNA sequencing (scRNA-seq) data from three different tissues with varying numbers of sub-cell types. **b**. To identify sub-cell types from scRNA-seq data, optimal clustering methods, such as ChooseR and multiK, have been developed. These methods begin with a dimensionality reduction (DR) step. This DR process starts with data preprocessing, such as log-normalization, to address bias and skewness in the scRNA-seq data distribution. Optionally, feature selection is used to filter out genes with low variance. Then, the dimension of this scaled data is reduced by applying principal component analysis (PCA). During this step, conventional optimal clustering methods typically recommend using the default number of principal components (PCs). As a result, all three datasets are reduced to identical dimensions despite having different numbers of cells or types of cells and, hence, different amounts of biological information. **c**. From these reduced datasets with identical dimensions, ChooseR and MultiK construct a graph, which consists of vertices of cells. To identify this graph’s optimal sub-communities (i.e., clusters), both methods repeatedly perform graph-based clustering to obtain multiple clustering results. Among the multiple clustering results, ChooseR and MultiK select the optimal clustering results using the co-clustering-based clustering robustness estimation and the clustering stability metric, reverse proportion of ambiguous clustering (rPAC), respectively. **d**. In these consensus clustering procedures, we replaced the conventional DR process with scLENS, a recently developed DR method that utilizes a data-driven signal selection approach. By employing noise filtering methods based on random matrix theory and a signal robustness test, scLENS adaptively determines the number of dimensions according to the amount of biological information in the data. This adaptive feature of scLENS enables ChooseR and MultiK to construct graphs that more effectively capture the biological patterns from various datasets with diverse amounts of biological information, potentially enhancing the quality of clustering results. For a comprehensive evaluation, we benchmarked our pipeline against both the original algorithms and three alternative dimensionality reduction (DR) approaches. The first is the default PCA setting in Scanpy(18), which uses a fixed number of 50 PCs (*N*=50 in b). **e**. The second alternative uses the elbow method(24), which selects the optimal number of PCs for PCA by identifying the point of diminishing returns in explained variance (orange circle in Fig. 1e). The third is scVI(19), a deep generative model that uses a variational autoencoder to learn a non-linear latent representation of the data as an alternative to PCA.

By replacing these DR steps with scLENS, we developed a fully data-driven clustering pipeline (Fig. 1c and 1d). To comprehensively evaluate this new approach, we benchmarked its performance against the original algorithms and three alternative methods. These methods were derived from the original algorithms by replacing the conventional DR with one of the following: PCA with Scanpy’s default settings [18] (Fig. 1b, *N* = 50), PCA using the elbow method [24] for automatic PC selection, and VAE-based tool scVI [19] (Fig. 1e). Across 33 datasets containing ground-truth cell-type annotations, our results showed overall improvements not only over the original MultiK and ChooseR but also over the alternative DR approaches using the elbow method and scVI. These findings confirm that integrating scLENS with optimal clustering algorithms enables more precise classification of cell populations, which, in turn, is expected to benefit downstream analyses, such as DEA analysis, trajectory analysis, and pseudotime analysis.

## Methods

### Quality Control (QC)

All analyses were performed on datasets that had undergone quality control. Specifically, cells with fewer than 200 expressed genes and genes expressed in fewer than 15 cells were filtered out. In addition, cells with a mitochondrial gene proportion exceeding 5% were removed. These steps ensured that only high-quality cells were retained for downstream analyses.

### Data Preprocessing

Following QC, data preprocessing was performed according to the specific requirements of each analytical method. For standard downstream analyses, the filtered count matrix was processed through a pipeline involving size factor normalization to correct for sequencing depth, a log transformation (*log*(*x*+1)) to stabilize variance, and z-score scaling to normalize each gene’s mean to zero and standard deviation to one (Fig. 1b).

After z-score scaling, scLENS performs an additional L2-normalization step as its final preprocessing stage to correct signal distortion [27] (Fig. 1d). In contrast, the scVI model was applied directly to the raw count matrix after the initial QC filtering (Fig. 1e). This is because scVI performs normalization internally as part of its generative modeling process and does not require the aforementioned transformation and scaling steps [19].

### Clustering Algorithms

- ChooseR [29]: Full clustering is initially performed on the preprocessed dataset to establish reference labels at a given resolution. Next, after applying PCA to the entire dataset, 100 clustering runs are conducted at this resolution on random 80% subsamples extracted from the resulting 100-dimensional embedding (Fig. 1b, *N* = 100). From each run, a pairwise co-clustering matrix is computed, with each entry representing the frequency with which two cells are grouped together (Fig. 1c). This co-clustering matrix is then converted into a distance matrix by subtracting each co-clustering frequency from 1, and per-cell silhouette scores are calculated from these distances. The scores of the cells within the same clusters are then averaged to obtain the cluster-level silhouette scores, indicating the robustness of each cluster. This entire process is repeated over a set of ten resolution parameters (resolution parameter = [0.3, 0.5, 0.8, 1, 1.2, 1.6, 2, 4, 6, 8]). For each resolution, the 95% confidence interval (CI) for the distribution of cluster silhouette scores is calculated. After first identifying the 95% CI with the highest lower bound, the near-optimal clustering parameter is selected as the one that yields the most clusters whose median silhouette scores fall within this interval.
- MultiK [30]: First, 100 subsamples are generated by randomly selecting 80% cells from the preprocessed data. PCA is then applied to each subsample to produce a 30-dimensional embedding (Fig. 1b, *N* = 30). These 100 PCA embeddings are subsequently used for clustering via the standard Seurat workflow across 40 resolution parameters (ranging from 0.05 to 2.0 with a step size of 0.05), resulting in 4,000 candidate clustering results (Fig. 1c). For each candidate clustering result, a consensus matrix is constructed to quantify the frequency with which cell pairs co-cluster. From the cumulative distribution function (CDF) curve of this matrix, the relative proportion of ambiguous clustering (rPAC) score is calculated to assess the stability for each number of clusters (*k*), using the following formula:

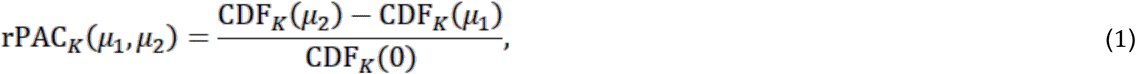

where *μ*_1_ = 0.1, *μ*_1_ = 0.9. Finally, the optimal clustering result is chosen based on two criteria: an rPAC score close to zero, which indicates robust clustering, and a high frequency of occurrence for that *k*.

### Integration with scLENS

Originally, scLENS was implemented in Julia, while ChooseR and MultiK were developed as R packages [27]. To facilitate the integration of scLENS with ChooseR and MultiK, all three packages were converted into Python implementations. Performance tests comparing the converted packages with their original versions confirmed that the default settings yielded equivalent performance. Once equivalence was verified, scLENS was integrated into the clustering workflows to enable a fully automated, data-driven selection of PCs (Fig. 1d). Specifically, for ChooseR, the conventional PCA step was replaced by scLENS in a single run. On the other hand, for MultiK—which typically performs multiple PCA runs across subsamples—scLENS was applied to the first subsampled dataset to determine the optimal number of PCs, and this determined value was then reused in subsequent subsampling iterations to enhance computational efficiency.

### Alternative DR methods

- PCA with Scanpy’s default settings [18]: First, the raw count data is preprocessed by removing its size factors and applying a logarithmic transformation, following the standard Scanpy pipeline. Subsequently, PCA is applied to this data to obtain the default 50 PCs (Fig. 1b, *N* = 50).
- scVI [19]: After QC, the scVI model is built from the raw count data using its default parameters (Fig. 1e). Key parameters include a zero-inflated negative binomial distribution for gene likelihood and a hidden layer size of 128 units. The model is then trained for 400 epochs to obtain a 10-dimensional latent space from the final model (Fig. 1e).
- PCA with the Elbow Method: First, the raw count data is preprocessed through size-factor removal, logarithmic transformation, and Z-score scaling[18] (Fig. 1e). PCA is then performed on the resulting scaled data. Finally, we use the kneed Python package [31] to identify the elbow point in the explained variance curve, which indicates the optimal number of PCs (orange circle in Fig. 1e).

## Results

### scLENS integration increases the Clustering Performance of ChooseR and MultiK

To benchmark the impact of different DR methods on consensus clustering, we assessed how the clustering performance of the two consensus clustering packages, ChooseR and MultiK, varied when provided with embeddings from: (1) scLENS, (2) PCA using Scanpy’s default settings, (3) PCA with the number of components selected by the elbow method, and (4) scVI (Fig. 1b, 1d, and 1e) [18,19,24,27]. The performance of these combinations was assessed on 33 real and simulated datasets derived from three sources: ZhengMix, a PBMC blood cell dataset; Tabula Muris, a comprehensive mouse tissue dataset; and a T-cell dataset obtained from the Cross-Tissue Immune Cell Atlas, representing immune cell data [32–34]. These datasets encompass varying levels of sparsity, skewness in data distributions, and population sizes, which can significantly influence clustering performance (Table 1).

**Table 1.** To evaluate the effectiveness of replacing conventional dimensionality reduction (DR) methods with scLENS in terms of clustering performance, we used 33 datasets with varying amounts of information influenced by factors such as the number of cells, cell types, and the sparsity level.

| Cat. | Name | Ncells | Ngenes | Sparsity | CV(TGC) | Avg.<br>Cluster size | Imbalanced<br>ratio | # of<br>signals |
| --- | --- | --- | --- | --- | --- | --- | --- | --- |
| sim_Tcell | 2250-1220 | 2951 | 10652 | 0.91 | 0.52 | 230.77 | 53.00 | 20 |
| sim_Tcell | 2251-1179 | 3000 | 10975 | 0.91 | 0.52 | 230.77 | 71.50 | 19 |
| sim_Tcell | 2972-1523 | 3000 | 11361 | 0.89 | 0.51 | 230.77 | 85.78 | 20 |
| sim_Tcell | 3276-1042 | 3000 | 11518 | 0.88 | 0.32 | 230.77 | 119.50 | 22 |
| sim_Tcell | 3689-1878 | 3000 | 11622 | 0.88 | 0.51 | 230.77 | 81.89 | 19 |
| sim_Tcell | 4249-2916 | 2971 | 11600 | 0.87 | 0.68 | 230.77 | 79.88 | 28 |
| sim_Tcell | 4320-2134 | 3000 | 11818 | 0.86 | 0.49 | 230.77 | 81.33 | 22 |
| sim_Tcell | 4797-2332 | 3000 | 11942 | 0.86 | 0.49 | 230.77 | 71.10 | 21 |
| sim_Tcell | 5930-3210 | 2988 | 12048 | 0.85 | 0.54 | 230.77 | 63.27 | 26 |
| sim_Tcell | 5960-1400 | 3000 | 12086 | 0.84 | 0.24 | 230.77 | 81.56 | 21 |
| sim_Tcell | 5974-3858 | 2979 | 12050 | 0.85 | 0.64 | 230.77 | 52.15 | 29 |
| sim_Tcell | 6006-2261 | 2998 | 12072 | 0.84 | 0.38 | 230.77 | 102.71 | 24 |
| sim_Tcell | 6083-4496 | 2974 | 12086 | 0.85 | 0.73 | 230.77 | 73.11 | 31 |
| Tabula_Muris | T_muris_1019 | 1019 | 12199 | 0.82 | 0.61 | 101.90 | 1.21 | 36 |
| Tabula_Muris | T_muris_2006 | 2006 | 12812 | 0.83 | 0.68 | 200.60 | 1.19 | 49 |
| Tabula_Muris | T_muris_3016 | 3016 | 13355 | 0.84 | 0.73 | 301.60 | 1.19 | 67 |
| Tabula_Muris | T_muris_3966 | 3966 | 13556 | 0.85 | 0.67 | 396.60 | 1.19 | 75 |
| Tabula_Muris | T_muris_5033 | 5033 | 13949 | 0.85 | 0.64 | 503.30 | 1.17 | 93 |
| Tabula_Muris | tmp_1 | 4827 | 11355 | 0.83 | 0.35 | 689.57 | 18.33 | 150 |
| Tabula_Muris | tmp_2 | 3501 | 12117 | 0.84 | 0.45 | 583.50 | 17.53 | 218 |
| Tabula_Muris | tmp_3 | 4244 | 13107 | 0.79 | 0.66 | 606.29 | 12.70 | 228 |
| Tabula_Muris | tmp_4 | 4313 | 13540 | 0.84 | 0.59 | 431.30 | 8.28 | 127 |
| Tabula_Muris | tmp_5 | 2485 | 10294 | 0.8 | 0.47 | 621.25 | 27.17 | 139 |
| ZhengMix | z_data_12073 | 11971 | 11850 | 0.95 | 0.46 | 1468.75 | 1.21 | 23 |
| ZhengMix | z_data_2410 | 2396 | 8194 | 0.93 | 0.45 | 301.25 | 1.18 | 10 |
| ZhengMix | z_data_3308 | 3288 | 9057 | 0.94 | 0.43 | 413.50 | 2.05 | 9 |
| ZhengMix | z_data_3706 | 3668 | 9581 | 0.94 | 0.44 | 463.25 | 3.44 | 15 |
| ZhengMix | z_data_3869 | 3838 | 9501 | 0.94 | 0.44 | 483.63 | 1.15 | 13 |
| ZhengMix | z_data_4292 | 4254 | 9813 | 0.94 | 0.46 | 536.50 | 1.97 | 12 |
| ZhengMix | z_data_4757 | 4721 | 9962 | 0.94 | 0.45 | 586.25 | 1.40 | 13 |
| ZhengMix | z_data_5730 | 5683 | 10322 | 0.95 | 0.48 | 716.25 | 12.71 | 14 |
| ZhengMix | z_data_785 | 778 | 4803 | 0.89 | 0.43 | 98.13 | 1.16 | 10 |
| ZhengMix | z_data_7993 | 7941 | 11019 | 0.95 | 0.45 | 975.88 | 1.28 | 19 |

Across these 33 datasets, scLENS integration yielded the most substantial improvements in the clustering performance. When integrated with scLENS, clustering performance, measured by Element-Centric Similarity (ECS) [35] against true labels, increased by an average of ∼ 14% for ChooseR and ∼ 10% for MultiK (Fig. 2a and 2b). In contrast, the alternative methods provided only slight improvements and even caused considerable performance degradation in some instances. For example, integrating ChooseR with Scanpy’s PCA or scVI resulted in a performance gain of merely ∼ 6% or less (green and pink bars in Fig. 2a), which was comparable to the baseline performance of the original packages. The elbow method for PC selection was even less effective, decreasing the average performance of ChooseR by ∼37% and showing the worst results for MultiK (Fig. 2a, cyan bar). For MultiK, both Scanpy’s PCA and scVI also showed lower performance compared to the original package (Fig. 2b). These results underscore the unique effectiveness of scLENS in enhancing clustering accuracy.

**Figure 2.**
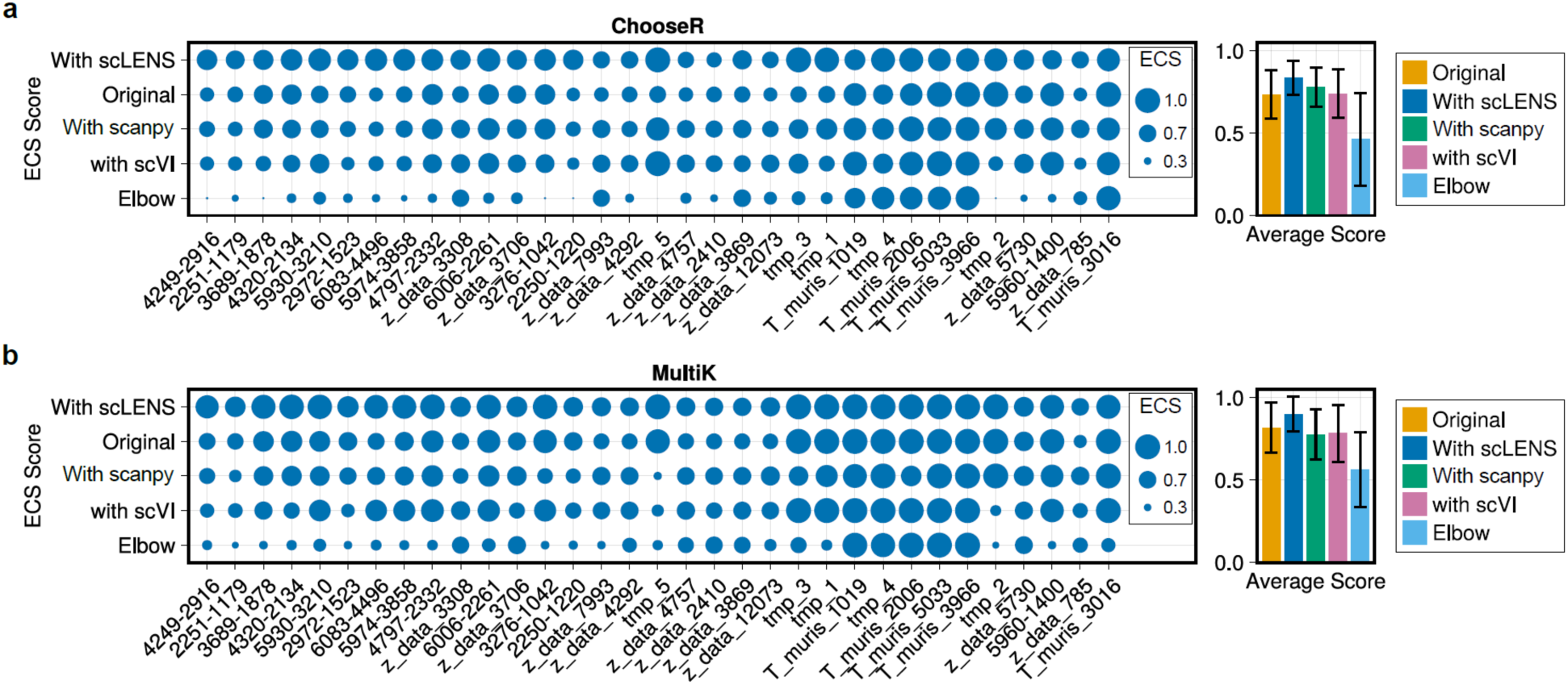
Integrating conventional clustering packages with scLENS enhances clustering output. **a-b**. The integration of scLENS significantly enhances the clustering performance of both ChooseR and MultiK, as measured by the element-centric similarity (ECS). Specifically, it substantially improves the performance of ChooseR by ∼14% (a, blue bar) compared to that of the original package (a, yellow bar). In contrast, integrating Scanpy’s PCA or scVI offers merely comparable performance to that of the original ChooseR (a, green and pink bars). Furthermore, using the elbow method for PC selection severely degrades ChooseR’s performance by ∼37% (a, cyan bar). Similarly, while alternative DR methods cause MultiK to underperform relative to its baseline (b, green, pink, and cyan bars), scLENS substantially improves its clustering performance by ∼10% (b, blue bar).

### The improved embedding quality from scLENS drives the enhancement in clustering performance of ChooseR and MultiK

While these average results highlighted the overall superiority of scLENS, a closer analysis revealed that the effectiveness of scLENS integration was highly dependent on the dataset (Fig. 3a). Specifically, it produced substantial performance gains for both ChooseR and MultiK on the simulated T-cell and real ZhengMix datasets, with increases ranging from ∼14% to ∼18% over their respective baselines (Fig. 3a). The simulated Tabula Muris data presented a unique case: scLENS still improved ChooseR’s performance by ∼9%, while MultiK’s performance remained at its optimal baseline (ECS ≈ 1), precluding further gains (Fig. 3a).

**Figure 3.**
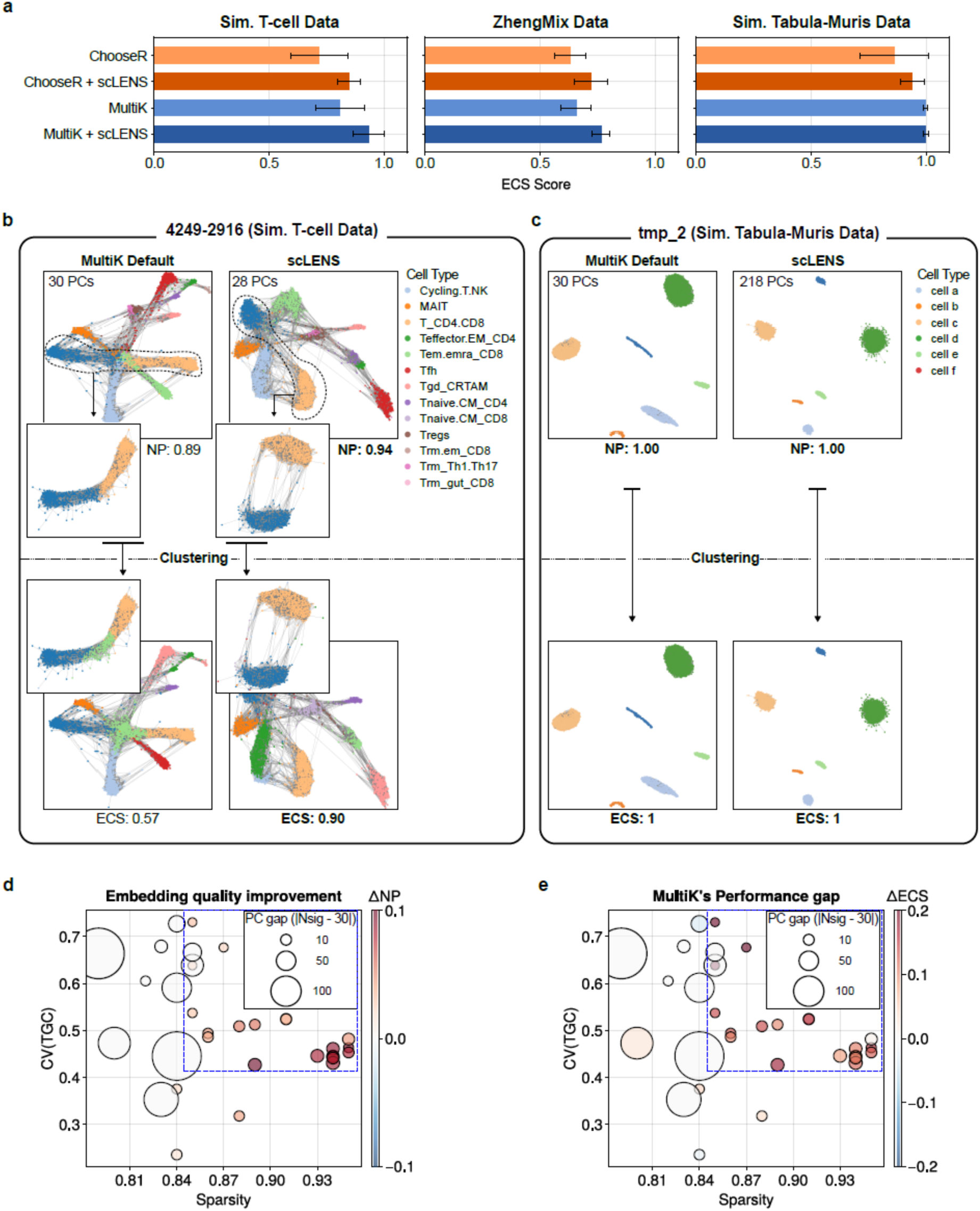
Clustering performance improvements from applying scLENS vary depending on data properties. **a**. The effectiveness of scLENS is highly dependent on the data type. scLENS integration yields substantial performance gains in clustering for both packages on T-cell and ZhengMix data (dark blue and orange bars). On Tabula-Muris data, scLENS integration improves ChooseR’s performance while MultiK’s performance remains at its maximum (ECS ∼ 1), having hit a ceiling. (dark blue bar in Sim. Tabula-Muris Data). **b**. The effectiveness of scLENS integration stems from its ability to generate higher-quality embedding. For instance, in the simulated T-cell dataset “4249-2916,” a graph constructed from the 28-dimensional embedding derived by scLENS has more edges within each cell type and fewer edges between cell types than the graph constructed from MultiK’s default 30-dimensional embedding. This change in connectivity creates a clearer cell type separation (sub-graphs with two cell types in small boxes). This clearer separation leads to a substantial increase in neighbor purity (NP)—a measure of how well neighboring cells share the same type. Higher NP indicates a higher-quality embedding, enabling MultiK to more accurately identify cell types and thereby enhance clustering performance (bottom panels). **c**. In contrast, such performance gains are not observed in one of the simulated Tabula-Muris datasets, “tmp_3.” In that dataset, only marginal improvements in NP (i.e., embedding quality) appear, despite a large gap between the 218 detected signals and the default of 30, leading to no meaningful increase in overall clustering performance. **d-e.** The embedding quality difference (d) and clustering performance difference (e) between MultiK with default settings and MultiK integrated with scLENS are quantified by NP and ECS, respectively. Furthermore, the size of the circles indicates the absolute difference (PC gap) between the dimension scLENS determines (based on its detected signals) and the fixed default PC dimension of 30. The degree of improvement in graph quality afforded by scLENS - depends on the sparsity level and coefficient of variation (CV) of total gene expression (TGC). For instance, in datasets with low sparsity or CV(TGC) (outside of the dashed box), scLENS and MultiK’s original embedding lead to graphs with similar NP (a), and thus, their performances are similar (b) although different numbers of signals are used (PC gap). On the other hand, when CV(TGC) ≥ 0.4 and Sparsity level ≥ 0.845 (inside of the dashed box), MultiK integrated with scLENS demonstrates more significant embedding-quality improvements in terms of NP as the PC gap grows (inside the blue dashed box in d). This substantial improvement in embedding quality, in turn, results in better clustering performance for most cases (inside the blue dashed box in e).

These performance variations across different data types stem from differences in the quality of embedding enhancement achieved by scLENS. For example, in one of the simulated T cell datasets, “4249-2916,” compared to the graph constructed on the default 30-PC embedding, the graph built on the 28-PC embedding derived by scLENS showed substantially higher neighbor purity (NP) [36]—defined as the proportion of neighboring nodes sharing the same cell type as a given node (top panels in Fig. 3b). This elevated NP, indicating more cohesive clusters of cells belonging to the same cell type, led to better cluster boundaries and subsequently enhanced the clustering performance for MultiK (bottom panels in Fig. 3b). However, this performance enhancement was not observed in one of the simulated Tabula-Muris datasets, “tmp_3,” despite a substantial difference between the number of PCs detected by scLENS and the default 30 PCs, as the NP increase remained minimal (Fig. 3c).

### Easy vs. Difficult Problems: How data variability and sparsity levels affect clustering performance

We further investigated why the degree of improvement from scLENS varied, especially in MultiK, by examining embedding quality in terms of NP under different sparsity levels and across inter-cell variability in total gene expression—quantified as the coefficient of variation of total gene expression (CV(TGC)) [27]. This embedding quality, quantified via NP differences between default and scLENS-driven embeddings, tended to show limited gains in data with low sparsity (≤0.84) or low CV(TGC) (≤0.4), even when the PC gap was substantial (big circles outside the blue dashed box in Fig. 3a). Such limited gains suggest that these “easier” datasets, characterized by abundant information due to low sparsity and minimal distortions from lower skewness in data distribution, are adequately handled by default preprocessing, leaving minimal scope for further improvement. In contrast, for “difficult” datasets with greater sparsity (>0.84) and CV(TGC) (>0.4), a larger PC gap can lead to greater NP gains (circles within the blue dashed box in Fig. 3d). These NP gains, in turn, often result in improved clustering performance as measured by ECS (Fig. 3e).

## Discussion

In general, DR and clustering are essential steps in scRNA-seq analysis; however, determining the optimal number of reduced dimensions and clusters still relies heavily on subjective user input, which can undermine the reliability of the results [17,18,20]. To address this issue, automated techniques, including ChooseR and MultiK, have been developed [29,30]. However, these methods fix the number of PCs during the DR process, making it difficult to consider them entirely data-driven. Therefore, we proposed a data-driven approach integrating scLENS [27] into these algorithms, significantly reducing user intervention from dimension reduction to clustering in scRNA-seq data analysis.

This combined algorithm was effective in enhancing clustering accuracy when applied to 33 real and simulated scRNA-seq datasets (Fig. 2). Specifically, compared with the default settings, ChooseR exhibited an improvement of ∼ 14 % in performance, and MultiK ∼ 10 % (blue bars in Fig. 2). In particular, these improvements were most pronounced in challenging datasets, where high CV(TGC) can lead to signal distortion, and greater sparsity indicates less information (Fig. 3d and 3e).

This significant performance gain becomes even more evident when contrasted with other widely used DR approaches. For instance, while the elbow method offers the convenience of automatically selecting PCs, it significantly degrades the clustering performance of ChooseR and MultiK (cyan bars in Fig. 2). Furthermore, other widely used DR methods, such as scVI and Scanpy’s PCA, failed to deliver significant improvements, offering performance that was merely comparable to the original packages (green and pink bars in Fig. 2). These results confirm that the substantial performance gains are uniquely attributable to the integration of scLENS, an advantage not replicated by the other methods.

Despite these clear performance advantages, the algorithm’s comprehensive evaluation process introduces scalability issues, as both memory usage and computation time increase rapidly with large-scale data. Specifically, scLENS requires the computation of the complete set of eigenvalues and eigenvectors, which significantly increases its memory requirements and limits its speed when handling massive datasets [27]. Similarly, as the number of cells increases, the repeated clustering required by MultiK or ChooseR leads to rapid increases in computation time and memory consumption as well [29,30]. To mitigate these scalability issues, the clustering algorithm employing MultiK and ChooseR can be replaced with the recently developed GVE-Leiden clustering algorithm, which leverages parallel computing to improve clustering speed significantly [37]. Furthermore, the speed and memory efficiency of scLENS may be enhanced in future research through approximate spectral analysis [38].

In this study, we demonstrated that replacing the conventional DR process, which relies on a fixed number of PCs, with the scLENS data-driven signal selection approach improves performance by 10 - 15%. Given that most downstream analyses depend on low-dimensional embedding and clustering results, combining scLENS with conventional algorithms is expected to enhance the performance of many existing methods for scRNA-seq data analysis. For instance, a recently developed package for evaluating clustering consistency has proven both highly efficient and effective in estimating clustering stability by successfully utilizing scLENS for its DR step [39].

Furthermore, this approach is applicable not only to scRNA-seq but also to various omics data, such as single-cell chromatin accessibility sequencing and spatial transcriptomics, which similarly require dimension reduction and clustering analysis [40–42]. Additionally, many analysis tools employing nonlinear neural network models—such as autoencoders or VAEs— have been developed for the dimension reduction step in extended data analyses [12,19]. To properly evaluate the performance of these tools, rather than simply comparing them with PCA results obtained using a fixed dimension, one can compare them with PCA results optimally selected via scLENS, thereby determining the extent to which nonlinear models can overcome the limitations of linear methods. In summary, the highly extensible Python version of the scLENS algorithm proposed in this study is expected to be widely adopted in various downstream analyses in the future, owing to its high user convenience and performance.

## Data availability

The 33 real and simulated datasets used in this study were derived from three sources:

- Zheng datasets [33]: Ten real datasets representing PBMC blood cell data were obtained from the Zheng datasets provided by 10x Genomics (https://www.10xgenomics.com/resources/datasets). These datasets exhibit a wide range of cluster sizes and ratios, thereby providing diverse clustering scenarios.
- Simulated Tabula Muris dataset [32]: Between 4 and 10 distinct cell types were randomly selected from various tissue types within the Tabula Muris dataset (https://tabula-muris.ds.czbiohub.org/) to minimize inter-cell type correlations. Using scDesign2 [43], we trained a model on these selected cells to generate simulated datasets with variable cluster sizes and imbalanced ratios, thereby providing diverse clustering scenarios.
- Simulated T-cell datasets [34]: T-cell data were extracted from the immune cell dataset available from the Cross-Tissue Immune Cell Atlas (https://www.tissueimmunecellatlas.org). Subsequently, using scDesign2, we generated multiple simulated T-cell datasets by varying sequencing depth and data skewness. This procedure produced datasets with different levels of quality, as reflected in metrics such as sparsity and the coefficient of variation of total gene expression (CV(TGC)).

In addition to the 33 datasets used in this study, the Python packages for scLENS, multiK, and ChooseR are publicly available at https://github.com/Mathbiomed/scLENSpy. (the repository URL will be disclosed upon acceptance; until then, the code is provided as supplementary material).

## Code Availability

A fully automated clustering package, integrating the MultiK and ChooseR optimal clustering algorithms with the scLENS data-driven dimensionality reduction (DR) method, is freely available on GitHub: https://github.com/Mathbiomed/scLENSpy.

## Ethics Statement

This study used publicly available and simulated datasets. All data were previously published and did not involve any new experiments on human subjects. Therefore, this research did not require an ethics statement or approval from an institutional review board.

## Acknowledgments.

This work was supported by the Institute for Basic Science IBS-R029-C3 (to J.K. Kim (Jae Kyoung))

## References

1. Macosko EZ, Basu A, Satija R, Nemesh J, Shekhar K, Goldman M, et al. Highly Parallel Genome-wide Expression Profiling of Individual Cells Using Nanoliter Droplets. Cell. 2015;161:1202–14.

2. Luecken MD, Theis FJ. Current best practices in single-cell RNA-seq analysis: a tutorial. Molecular Systems Biology. 2019;15:e8746.

3. Kharchenko PV. The triumphs and limitations of computational methods for scRNA-seq. Nat Methods. 2021;18:723–32.

4. Andrews TS, Kiselev VY, McCarthy D, Hemberg M. Tutorial: guidelines for the computational analysis of single-cell RNA sequencing data. Nat Protoc. 2021;16:1–9.

5. Ahlmann-Eltze C, Huber W. Comparison of transformations for single-cell RNA-seq data. Nat Methods. 2023;20:665–72.

6. Kiselev VY, Andrews TS, Hemberg M. Challenges in unsupervised clustering of single-cell RNA-seq data. Nat Rev Genet. 2019;20:273–82.

7. Finak G, McDavid A, Yajima M, Deng J, Gersuk V, Shalek AK, et al. MAST: a flexible statistical framework for assessing transcriptional changes and characterizing heterogeneity in single-cell RNA sequencing data. Genome Biol. 2015;16:278.

8. Trapnell C. Defining cell types and states with single-cell genomics. Genome Res. 2015;25:1491–8.

9. Trapnell C, Cacchiarelli D, Grimsby J, Pokharel P, Li S, Morse M, et al. The dynamics and regulators of cell fate decisions are revealed by pseudotemporal ordering of single cells. Nat Biotechnol. 2014;32:381–6.

10. Bergen V, Lange M, Peidli S, Wolf FA, Theis FJ. Generalizing RNA velocity to transient cell states through dynamical modeling. Nat Biotechnol. 2020;38:1408–14.

11. Van Den Berge K, Roux De Bézieux H, Street K, Saelens W, Cannoodt R, Saeys Y, et al. Trajectory-based differential expression analysis for single-cell sequencing data. Nat Commun. 2020;11:1201.

12. Tran D, Nguyen H, Tran B, La Vecchia C, Luu HN, Nguyen T. Fast and precise single-cell data analysis using a hierarchical autoencoder. Nat Commun. 2021;12:1029.

13. Abdi H, Williams LJ. Principal component analysis. WIREs Computational Stats. 2010;2:433–59.

14. Becht E, McInnes L, Healy J, Dutertre C-A, Kwok IWH, Ng LG, et al. Dimensionality reduction for visualizing single-cell data using UMAP. Nat Biotechnol. 2019;37:38–44.

15. McInnes L, Healy J, Saul N, Großberger L. UMAP: Uniform Manifold Approximation and Projection. JOSS. 2018;3:861.

16. Kobak D, Berens P. The art of using t-SNE for single-cell transcriptomics. Nat Commun. 2019;10:5416.

17. Hao Y, Hao S, Andersen-Nissen E, Mauck WM, Zheng S, Butler A, et al. Integrated analysis of multimodal single-cell data. Cell. 2021;184:3573–3587.e29.

18. Wolf FA, Angerer P, Theis FJ. SCANPY: large-scale single-cell gene expression data analysis. Genome Biol. 2018;19:15.

19. Lopez R, Regier J, Cole MB, Jordan MI, Yosef N. Deep generative modeling for single-cell transcriptomics. Nat Methods. 2018;15:1053–8.

20. Elhaik E. Principal Component Analyses (PCA)-based findings in population genetic studies are highly biased and must be reevaluated. Sci Rep. 2022;12:14683.

21. Mima A, Kimura A, Ito R, Hatano Y, Tsujimoto H, Mae S-I, et al. Mechanistic elucidation of human pancreatic acinar development using single-cell transcriptome analysis on a human iPSC differentiation model. Sci Rep. 2025;15:4668.

22. Martins-Ferreira R, Calafell-Segura J, Leal B, Rodríguez-Ubreva J, Martínez-Saez E, Mereu E, et al. The Human Microglia Atlas (HuMicA) unravels changes in disease-associated microglia subsets across neurodegenerative conditions. Nat Commun. 2025;16:739.

23. Zhong H, Han W, Gomez-Cabrero D, Tegner J, Gao X, Cui G, et al. Benchmarking cross-species single-cell RNA-seq data integration methods: towards a cell type tree of life. Nucleic Acids Research. 2025;53:gkae1316.

24. Zhuang H, Wang H, Ji Z. findPC: An R package to automatically select the number of principal components in single-cell analysis. Martelli PL, editor. Bioinformatics. 2022;38:2949–51.

25. Horn JL. A Rationale and Test for the Number of Factors in Factor Analysis. Psychometrika. 1965;30:179–85.

26. Jolliffe IT. Principal component analysis. 2nd ed. New York: Springer; 2002.

27. Kim H, Chang W, Chae SJ, Park J-E, Seo M, Kim JK. scLENS: data-driven signal detection for unbiased scRNA-seq data analysis. Nat Commun. 2024;15:3575.

28. Virshup I, Bredikhin D, Heumos L, Palla G, Sturm G, Gayoso A, et al. The scverse project provides a computational ecosystem for single-cell omics data analysis. Nat Biotechnol. 2023;41:604–6.

29. Patterson-Cross RB, Levine AJ, Menon V. Selecting single cell clustering parameter values using subsampling-based robustness metrics. BMC Bioinformatics. 2021;22:39.

30. Liu S, Thennavan A, Garay JP, Marron JS, Perou CM. MultiK: an automated tool to determine optimal cluster numbers in single-cell RNA sequencing data. Genome Biol. 2021;22:232.

31. Satopaa V, Albrecht J, Irwin D, Raghavan B. Finding a “Kneedle” in a Haystack: Detecting Knee Points in System Behavior. 2011 31st International Conference on Distributed Computing Systems Workshops [Internet]. Minneapolis, MN, USA: IEEE; 2011 [cited 2025 July 1]. p. 166–71. Available from: http://ieeexplore.ieee.org/document/5961514/

32. The Tabula Muris Consortium, Overall coordination, Logistical coordination, Organ collection and processing, Library preparation and sequencing, Computational data analysis, et al. Single-cell transcriptomics of 20 mouse organs creates a Tabula Muris. Nature. 2018;562:367–72.

33. Zheng GXY, Terry JM, Belgrader P, Ryvkin P, Bent ZW, Wilson R, et al. Massively parallel digital transcriptional profiling of single cells. Nat Commun. 2017;8:14049.

34. Domínguez Conde C, Xu C, Jarvis LB, Rainbow DB, Wells SB, Gomes T, et al. Cross-tissue immune cell analysis reveals tissue-specific features in humans. Science. 2022;376:eabl5197.

35. Gates AJ, Wood IB, Hetrick WP, Ahn Y-Y. Element-centric clustering comparison unifies overlaps and hierarchy. Sci Rep. 2019;9:8574.

36. Cheng C-H, Chan C-P, Sheu Y-J. A novel purity-based k nearest neighbors imputation method and its application in financial distress prediction. Engineering Applications of Artificial Intelligence. 2019;81:283–99.

37. Sahu S. GVE-Leiden: Fast Leiden Algorithm for Community Detection in Shared Memory Setting [Internet]. arXiv; 2023 [cited 2025 Feb 10]. Available from: https://arxiv.org/abs/2312.13936

38. Lin L, Saad Y, Yang C. Approximating Spectral Densities of Large Matrices. SIAM Rev. 2016;58:34–65.

39. Kim H, Park I, Park J-E, Kim JK, Seo M, Kim JK. scICE: enhancing clustering reliability and efficiency of scRNA-seq data with multi-cluster label consistency evaluation. Nat Commun. 2025;16:6031.

40. Rodriques SG, Stickels RR, Goeva A, Martin CA, Murray E, Vanderburg CR, et al. Slide-seq: A scalable technology for measuring genome-wide expression at high spatial resolution. Science. 2019;363:1463–7.

41. Cusanovich DA, Hill AJ, Aghamirzaie D, Daza RM, Pliner HA, Berletch JB, et al. A Single-Cell Atlas of In Vivo Mammalian Chromatin Accessibility. Cell. 2018;174:1309–1324.e18.

42. Ståhl PL, Salmén F, Vickovic S, Lundmark A, Navarro JF, Magnusson J, et al. Visualization and analysis of gene expression in tissue sections by spatial transcriptomics. Science. 2016;353:78– 82.

43. Sun T, Song D, Li WV, Li JJ. scDesign2: a transparent simulator that generates high-fidelity single-cell gene expression count data with gene correlations captured. Genome Biol. 2021;22:163.

